# Anoctamin-1 is induced by TGF-beta and contributes to lung myofibroblast differentiation

**DOI:** 10.1101/2023.06.07.544093

**Authors:** Eleanor B. Reed, Shaina Orbeta, Bernadette A. Miao, Albert Sitikov, Bohao Chen, Irena Levitan, Julian Solway, Gökhan M. Mutlu, Yun Fang, Alexander A. Mongin, Nickolai O. Dulin

## Abstract

Idiopathic pulmonary fibrosis (IPF) is a devastating disease characterized by progressive scarring of the lungs and resulting in deterioration in lung function. Transforming growth factor-beta (TGF-β) is one of the most established drivers of fibrotic processes. TGF-β promotes transformation of tissue fibroblasts to myofibroblasts, a key finding in the pathogenesis of pulmonary fibrosis. We report here that TGF-β robustly upregulates the expression of the calcium-activated chloride channel Anoctamin-1 (ANO1) in human lung fibroblasts (HLF) at mRNA and protein levels. ANO1 is readily detected in fibrotic areas of IPF lungs in the same area with smooth muscle alpha-actin (SMA)-positive myofibroblasts. TGF-β-induced myofibroblast differentiation (determined by the expression of SMA, collagen-1 and fibronectin) is significantly inhibited by a specific ANO1 inhibitor, T16A_inh_-A01, or by siRNA-mediated ANO1 knockdown. T16A_inh_-A01 and ANO1 siRNA attenuate pro-fibrotic TGF-β signaling, including activation of RhoA pathway and AKT, without affecting initial Smad2 phosphorylation. Mechanistically, TGF-β treatment of HLF results in a significant increase in intracellular chloride levels, which is prevented by T16A_inh_-A01 or by ANO1 knockdown. The downstream mechanism involves the chloride-sensing “with-no-lysine (K)” kinase (WNK1). WNK1 siRNA significantly attenuates TGF-β-induced myofibroblast differentiation and signaling (RhoA pathway and AKT), whereas the WNK1 kinase inhibitor WNK463 is largely ineffective. Together, these data demonstrate that (i) ANO1 is a TGF-β-inducible chloride channel that contributes to increased intracellular chloride concentration in response to TGF-β; and (ii) ANO1 mediates TGF-β-induced myofibroblast differentiation and fibrotic signaling in a manner dependent on WNK1 protein, but independent of WNK1 kinase activity.

**NEW & NOTEWORTHY:** This study describes a novel mechanism of differentiation of human lung fibroblasts (HLF) to myofibroblasts – the key process in the pathogenesis of pulmonary fibrosis. TGF-β drives the expression of calcium-activated chloride channel anoctmin-1 (ANO1) leading to an increase in intracellular levels of chloride. The latter recruits chloride-sensitive With-No-Lysine (K) kinase (WNK1) to activate pro-fibrotic RhoA and AKT signaling pathways, possibly through activation of mammalian target of rapamycin complex-2 (mTORC2), altogether promoting myofibroblast differentiation.

## INTRODUCTION

Idiopathic pulmonary fibrosis (IPF) is a progressive, fatal disease characterized by parenchymal fibrosis and structural distortion of the lungs. Age-adjusted mortality due to pulmonary fibrosis is increasing (1), and it poses a vexing clinical challenge given the lack of effective therapy. IPF is a disorder of abnormal wound healing (2, 3), wherein the initial trigger is injury to the alveolar epithelial cell, followed by a non-resolving wound-healing response (4-6). Migration of interstitial fibroblasts to sites of injury and their differentiation to myofibroblasts are thought to be critical processes in pathogenesis of pulmonary fibrosis. Myofibroblasts secrete extracellular matrix proteins and pro-fibrotic factors which perpetuate tissue remodeling and fibrosis (7, 8). Myofibroblasts are invariably found in histologic sections of human lung specimens from IPF patients and are thought to be a critical pathogenic mechanism responsible for the progressive nature of IPF.

Transforming Growth Factor-β (TGF-β) is the most well-established driver of myofibroblast differentiation (9-11). Production of this cytokine has been localized to areas of fibrosis in both experimental pulmonary fibrosis and human disease (9, 12, 13). Lung-targeted overexpression of TGF-β results in the development of lung fibrosis in animal models of IPF (11, 14). Conversely, inhibition of TGF-β, via soluble TGF-β receptors, can inhibit in vivo fibrogenesis (15, 16). TGF-β signaling acts through transmembrane-receptor serine/threonine kinases that phosphorylate Smad transcription factors (Smad2/3), leading to their heteromerization with a common mediator Smad4, nuclear translocation of the Smad2/3/4 complex, and activation of transcription of TGF-β target genes (7, 17, 18). We and others have established that signaling downstream of Smad-dependent gene transcription is critical for myofibroblast differentiation; this includes the TGF-β-induced RhoA/SRF pathway (19-21) and AKT activation (22-24).

Anoctamine-1 (ANO1), also known as TMEM16A, DOG1 (Discovered On Gastrointestinal stromal tumors 1), ORAOV2 (ORAl cancer OVerexpressed), and TAOS-2 (Tumor Amplified and Overexpressed) was initially found to be overexpressed in a number of cancer tissues and is thought to contribute to cancer cell growth and tumorigenesis (25, 26). Subsequently, ANO1 was identified as a calcium-activated chloride channel (27-29). Reported cellular functions of ANO1 include the control of cancer cell proliferation, cell survival and migration (25, 26), secretory function in the epithelia (including airways, intestines, salivary glands, renal tubules and sweat glands) (30), induction of electrical pacemaker activity of interstitial cells of Cajal in gastrointestinal smooth muscles (31), control of acute pain sensation, chronic pain and anxiety-related behaviors (32, 33), and contribution to contraction of airway and vascular smooth muscles (32). Through an unbiased microarray analysis, we identified ANO1 as one of the top transcripts induced by TGF-β in primary cultured human lung fibroblasts (HLF), which was not previously recognized. We therefore sought to confirm this finding by appropriate biochemical approaches and to examine the role of ANO1 in control of intracellular chloride homeostasis and myofibroblast differentiation.

## MATERIALS AND METHODS

### Primary culture of human lung fibroblasts (HLF)

Human lung fibroblasts (HLF) were isolated from human lungs rejected for transplantation through the Gift of Hope Organ and Tissue Donor Network. IPF-HLF were isolated from lungs of IPF patients shortly after their removal during lung transplantation at the University of Chicago under the IRB protocol #14514A. Human lung tissue samples were placed in DMEM with antibiotics. Lung tissue was minced to ∼1 mm^3^ pieces, washed, and plated on 10-cm plates in growth media containing DMEM supplemented with 10% FBS and antibiotics. The medium was changed twice a week. After approximately 2 weeks, the explanted and amplified fibroblasts were trypsinized, cleared from tissue pieces by sedimentation, and further amplified as passage 1. Unless indicated, cells were grown in 12-well plates at a density of 1×10^5^ cells per well in a growth media for 24 hours, starved in DMEM containing 0.1% FBS for 48 hours (48-hour starvation is important to ensure quiescence in all cells), and treated with desired drugs for various times as indicated in figure legends. Primary cultures were used from passage 3 to 8.

### Immunohistochemistry

Non-fibrotic human lungs were obtained from the Gift of Hope Organ and Tissue Donor Network. Fibrotic lungs from IPF patients were obtained shortly after lung transplantation at the University of Chicago under the IRB protocol #14514A. Lung tissue specimens were fixed in formalin, washed in 70% ethanol and paraffin embedded. Paraffin-embedded tissue was sectioned (5 µm) and stained with control IgG or ANO1 antibodies (Abcam # ab53212, 3 µg/ml) by the University of Chicago Immunohistochemistry Core Facility.

### Immunofluorescent microscopy

Paraffin-embedded IPF human lung specimens were sectioned (5 μm) by the University of Chicago Human Tissue Resource Center. Sections were deparaffinized with xylenes and rehydrated with ethanol. They were then permeabilized and blocked with 1% BSA with 0.1% Triton X-100 in PBS for 1 hour. Sections were next incubated with primary antibodies for ANO1 (Abcam Ab53212, 3 µg/ml) and smooth muscle actin (SMA, Sigma A2547, 1:100) overnight at 4°C. After washing with PBS, sections were incubated with secondary goat anti-rabbit IgG conjugated to Alexa Fluor 488 (Invitrogen A-11034, 1:400) and donkey anti-mouse IgG conjugated to Alexa Fluor 647 (Invitrogen A-31571, 1:400) at room temperature for 1 hour. After washing with PBS sections were mounted with mounting medium containing DAPI (Abcam Ab104139). Fluorescence images were acquired on a Zeiss 3i Marianas spinning disk confocal microscope with the 40x / NA 1.3 oil (Plan-Apochromat) objective using Slidebook acquisition software (University of Chicago Integrated Light Microscopy Core). Negative controls included the omission of primary antibodies.

### siRNA transfection

HLF were plated at a density of 0.8 x10^5^ cells per well (12-well plates, for western blotting), or 1.6 x10^5^ cells per well (6-well plates, for real time qPCR) and grown for 24 hours. Cells were then transfected with 10 nM negative control siRNA or ANO1 siRNA using Lipofectamine RNAiMAX Reagent (Thermo Fisher Scientific, Waltham, MA) according to the manufacturer’s instructions and kept in growth media for an additional 24 hours, followed by serum starvation in 0.1% FBS for 48 hours, and then treatment with TGF-β for desired times.

### Real-time qPCR

Direct-Zol RNA mini prep kit (Zymo Research, Irvin, CA) was used to isolate total RNA following the manufacturer’s protocol. RNA was random primed and reverse transcribed using iScript cDNA synthesis kit (Bio-Rad, Hercules, CA) according to the manufacturer’s protocol. Real-time PCR analysis was performed using iTaq SYBR Green supermix with ROX (Bio-Rad) in a MyIQ single-color real-time PCR detection system (Bio-Rad). The ANO1 primers were: GCAGAGAGGCCGAGTTTCTG (forward) and GCTCAGCCACTTTGGGCTG (reverse).

### Western blotting

Cells were lysed in a buffer containing 8 M deionized urea, 1% SDS, 10% glycerol, 60 mM Tris-HCl, pH 6.8, 0.02% pyronin Y, and 5% β-mercaptoethanol. Lysates were sonicated for 5 seconds. Samples were then subjected to polyacrylamide gel electrophoresis and Western blotting with desired primary antibodies and corresponding horseradish peroxidase (HRP)-conjugated secondary antibodies and then developed by chemiluminescence reaction. Digital chemiluminescent images below the saturation level were obtained with a LAS-4000 analyzer, and signal intensity was quantified using Multi Gauge software (Fujifilm, Valhalla, NY). Specificity of primary antibodies was validated based on the appropriate molecular weight of target proteins and by the loss of the immunoreactivity in siRNA-mediated knockdown experiments (ANO1, WNK1) (see Supplemental Figure S1, https://doi.org/10.6084/m9.figshare.24514975).

### Measurements of intracellular chloride

Intracellular chloride levels ([Cl⁻]_i_) were determined using steady-state ^36^Cl⁻ accumulation, which is proportional to [Cl⁻]_i_ and can be converted to cytosolic concentration values if the precise volume of intracellular water is known (34, 35). HLF were replated into 6-well plates, grown in DMEM-FBS, and subsequently differentiated and treated as described above. On the day of assay, cells were additionally pretreated with ion transport inhibitors (concentrations and timing are specified in Results). ^36^Cl⁻ accumulation was initiated by adding to each well 1/10 volume of low-FBS assay medium containing 0.5 µCi/ml Na^36^Cl and 10 mM HEPES (final concentrations). HEPES was supplemented to minimize alkalinization during temporary removal of cells from the CO_2_ incubator. ^36^Cl⁻ accumulation was carried out for 30 min at 37°C in the CO_2_ incubator. Radioisotope uptake was terminated by aspirating ^36^Cl⁻-containing medium and four immediate washes with 2 ml of ice-cold medium containing (in mM) 300 mannitol, 3 MgSO_4_, 10 HEPES (pH 7.4). After the final wash, HLF were lysed in 1 ml of solution containing 2% SDS plus 8 mM EDTA. Half of each cell lysate sample was used for [^36^Cl] detection, and the other half used to determine protein levels in individual wells. ^36^Cl⁻ in each sample was counted with a TriCarb 4910 scintillation counter (Perkin Elmer, Waltham, MA) after adding Ecoscint A scintillation cocktail (National Diagnostics, Atlanta, GA). The [Cl⁻]_i_ values were quantified by normalizing ^36^Cl⁻ counts to specific ^36^Cl⁻ activity and either protein levels or cell numbers in each individual well (see Results).

### Materials

TGF-β was from EMD Millipore (GF111). The following antibodies for Western blotting were from Millipore-Sigma: smooth muscle α-actin (A5228, 10,000×), β-actin (A5441, 10,000×), α-tubulin (T6074, 10,000×), and myosin light chain (M4401, 1,000×). Antibodies against human collagen-1A1 (sc-28657, 1,000×) and AKT (sc-8312, 1,000×) were from Santa Cruz Biotechnology. Antibodies against ANO1 (14476, 1,000×), Smad2 (L1603, 1,000×), phospho-Smad2-Ser465/467 (138D4, 1,000×), phospho-MLC-Ser19 (3671, 1,000×) and phospho-AKT-Ser473 (9271, 1,000×) were from Cell Signaling Technology. Secondary HRP-conjugated antibodies for western blotting (3,000×) were from Millipore Sigma (40-139-32ML - anti-rabbit IgG, 40-125-32ML - anti-mouse IgG). For immunohistochemistry and immunofluorescent microscopy, ANO1 antibodies were from Abcam (ab53212, 3 µg/ml final concentration), and SMA antibodies were from Millipore-Sigma (A5228, 100×). T16A_inh_-A01 inhibitor was from EMD Millipore (613551). SB-431542 was from Cayman Chemical (13031), bumetanide (B3023) and 4,4′-Diisothiocyano-2,2′-stilbenedisulfonic acid (DIDS, 309795) were from Millipore Sigma. The WNK inhibitor WNK463 (HY-100626HY) was from MedChemExpress. ANO1 siRNA (1027417), WNK1 siRNA (1027417) and AllStars negative control siRNA (1027281) were from Qiagen. Radiolabeled ^36^Cl (Na^36^Cl, 0.1 mCi/ml, 16 mCi/g Cl) was from American Radiolabeled Chemicals.

### Statistical Analysis

All data are presented as mean values ± SD. Normally distributed data were analyzed using Student t-test or one-way ANOVA with the Tukey honest significant difference test post hoc correction for multiple comparisons, as appropriate. Normalized-to-controls and non-normally distributed data were analyzed using Kruskal-Wallis non-parametric test followed by Dunn’s post hoc correction for multiple comparisons. For data sets presented in Fig. 4, 5, 6, 7, and 8 we performed additional preplanned analyses exploring the effect of ANO1 inhibitor and ANO1 siRNA in TGFβ-treated cells. The latter analysis was conducted using a one-population t-test with values compared to 100%. Values of p<0.05 were considered statistically significant. All statistical analyses were performed in Prism v. 9.5 (GraphPad Software).

## RESULTS

### Expression of ANO1 in human lung myofibroblasts in cell culture and in human lung fibrotic tissues

Treatment of primary cultured human lung fibroblasts (HLF) with TGF-β resulted in a robust accumulation of ANO1 expression at mRNA and protein levels, as assessed by real time qPCR and western blotting (Figs. 1A, 1B). This effect was mediated by TGF-β receptor II (TGFBRII), as it was inhibited by the specific TGFBRII kinase inhibitor, SB-431542 (Fig. 1C). Given that myofibroblasts are significant contributors to tissue scaring during fibrotic processes, we then assessed the expression of ANO1 in lung explants from patients with idiopathic pulmonary fibrosis (IPF) as compared to non-fibrotic lungs. As shown in figure 2A, ANO1 expression in non-fibrotic (NF) lungs was largely detected in smooth muscle. In the IPF lungs, ANO1 expression was clearly observed in fibrotic areas with a patchy appearance (Fig. 2A). Double immunofluorescent staining demonstrated specific ANO1 signal in IPF tissue, which was localized to the same areas as the SMA-expressing myofibroblasts (Fig. 2B).

**Figure 1.**
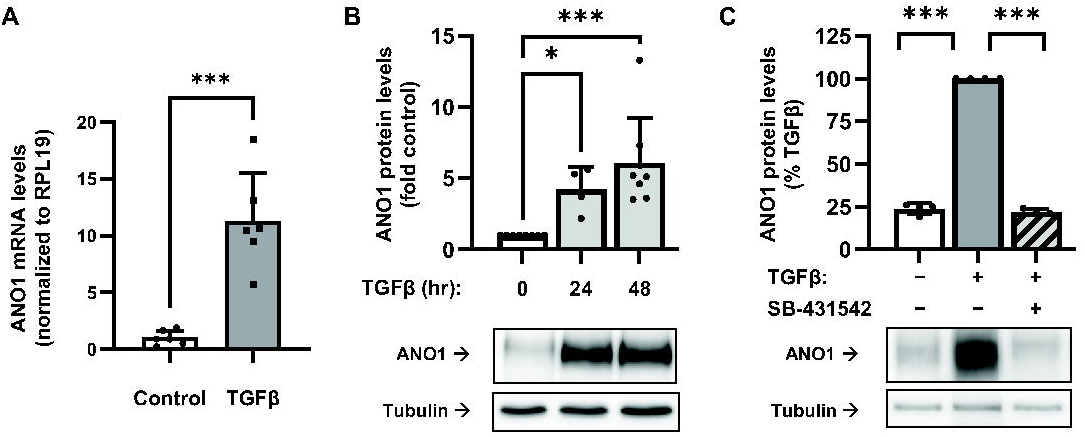
TGFβ-induced expression of anoctamin 1 (ANO1) in human lung fibroblasts. Human lung fibroblasts were serum starved in 0.1% FBS for 48 hr, followed by treatment with TGF-β (1 ng/ml) for 24 or 48 hrs, as indicated. **A,** ANO1 mRNA expression after 24-hr treatment with TGF-β as determined by RT-qPCR. Data are mean values ±SD. ***p<0.001, t-test. **B,** Western blot analysis and quantitation of ANO1 protein expression in response TGF-β. Data are mean values normalized to control cells ±SD. *, p<0.05, ***p<0.001, Kruskal-Wallis test with Dunn’s correction for multiple comparisons. **C,** Effect of the TGFBRII inhibitor, SB-431542 (5 µM, 30 minutes pre-incubation) on TGF-β-induced ANO1 protein expression (48 hrs). Shown are representative images of ANO1 immunoreactivity and quantification. Data are mean immunoreactivity values normalized to TGF-β-treated cells ±SD. ***p<0.001, one-population t-test with Bonferroni correction for multiple comparisons.

**Figure 2.**
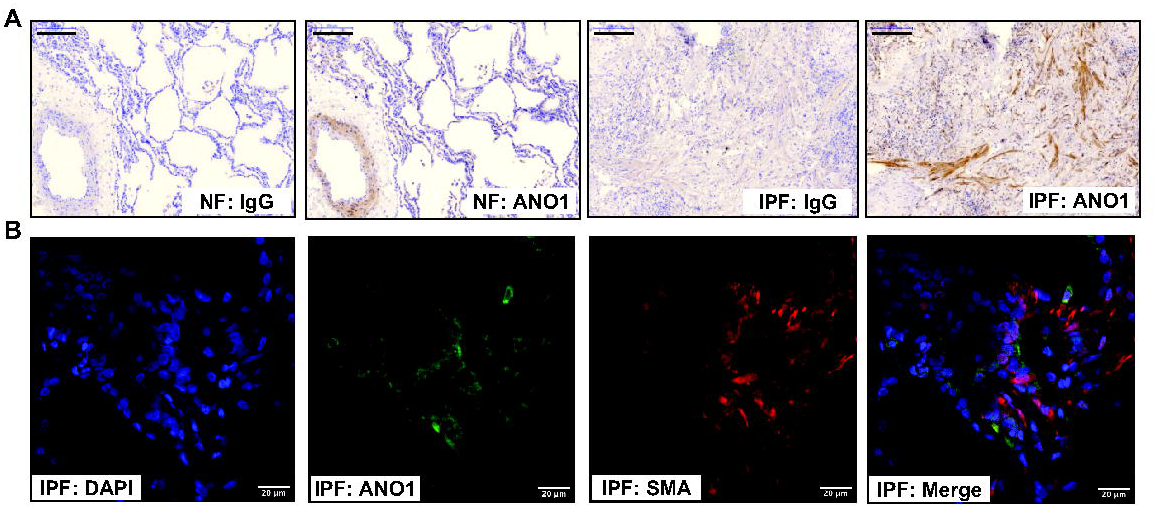
ANO1 protein expression in fibrotic areas of human lungs with idiopathic pulmonary fibrosis (IPF). **A,** Representative immunohistochemical images of non-fibrotic (NF) lungs and fibrotic areas of the IPF lungs. Sequential sections of the same lung were probed with either ANO1 antibody or IgG control. Scale bars = 200 μm. **B,** High magnification immunofluorescence images of section from an IPF lung, stained for ANO1 (green) and smooth muscle a-actin (SMA, red), and counterstained with DAPI (blue). Scale bars = 20 μm. Shown are representative images of lung sections from two IPF individuals. Note that we did not expect stains precisely overlapping because of membrane localization of ANO1 and cytosolic localization of ANO1.

### Effect of TGF-β on intracellular chloride levels, and the role of ANO1 in HLF

Given that ANO1 is a chloride channel, we next asked whether TGF-β affects intracellular chloride homeostasis in an ANO1-dependent manner. To assess intracellular chloride levels ([Cl^−^]_i_), we measured the steady state accumulation of ^36^Cl^−^ inside the cells. At equilibrium, ^36^Cl⁻ accumulation is directly proportional to [Cl^−^]_i_, and can be converted to cytosolic concentration values if the precise volume of intracellular water is known (34, 35). To determine appropriate conditions for measuring steady state ^36^Cl⁻ accumulation, we first assessed the kinetics of ^36^Cl⁻ uptake in HLF. As shown in Fig. 3A, intracellular ^36^Cl⁻levels reached equilibrium after approximately 10-15 min. The fast equilibration was determined by ^36^Cl⁻ uptake through several ion transport systems. Addition of bumetanide, an inhibitor of the Na^+^-K^+^-2Cl⁻ cotransporter (NKCC1), reduced steady-state chloride accumulation by ∼25% (n=4, p<0.05, data not shown). When bumetanide was combined with the anion exchanger blocker DIDS (4,4′-diisothiocyano-2,2′-stilbenedisulfonic acid), ^36^Cl⁻ uptake values were reduced by >90% and failed to saturate within 30 min (Fig. 3A). These results indicate that the main transport system that equilibrates ^36^Cl⁻ across the plasma membrane of HLFs is the DIDS-sensitive chloride-bicarbonate exchanger, with additional contributions from NKCC1 and possibly other DIDS-sensitive Cl⁻ channels.

**Figure 3.**
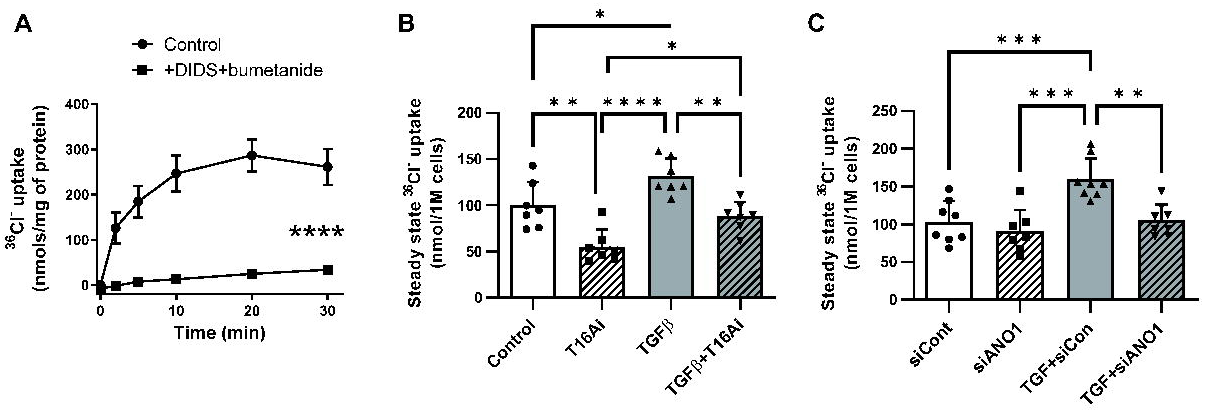
Effect of TGF-β and the role of ANO1 in control of intracellular Cl□ levels in human lung fibroblasts (HLF). **A,** Kinetics of ^36^Cl□ accumulation in HLF. Cl□ uptake was measured as radiotracer accumulation after adding ^36^Cl□ into cell culture medium. Serum-starved HLF cells were pretreated for 30 min with either vehicle or a combination of 1 mM DIDS plus 20 µM bumetanide. The same inhibitors were present in transport assay media. Data are mean values ±SD in three experiments. ****p<0.0001, repeated measures ANOVA; inhibitors vs. control. **B,** Effect of the ANO1 inhibitor, T16Ainh-A01 (T16Ai), on intracellular Cl□ levels in control and TGF-β-treated HLF. Intracellular Cl□ levels were measured as the steady state ^36^Cl□ accumulation and normalized to the number of cells per well as described in methods. T16Ainh-A01 (30 µM) was applied 1 hr before and was present during ^36^Cl□ transport assay. Data represent mean values ±SD from seven independent experiments in two HLF passages. *p<0.01, **p<0.01; ***p<0.001; One-way ANOVA with Tukey post hoc correction for multiple comparisons. **C,** The effect of siRNA-mediated ANO1 knockdown on intracellular Cl□ levels in control and TGFβ-treated primary human lung fibroblasts. HLF cells were transfected with the siRNA targeting ANO1 (siANO1) or the negative control RNA (siCont) for 24 hrs, serum starved for 48 hrs, and treated with TGF-β or vehicle control for 48 hrs. The steady-state ^36^Cl□ accumulation was measured and normalized as in panel B. Data represent mean values ±SD of eight independent experiments performed in two HLF passages. **p<0.01, ***p<0.001, One-way ANOVA with Tukey post hoc correction for multiple comparisons.

We next examined the effect of TGF-β and the pharmacological inhibitor of ANO1, T16A_inh_-A01 (36), on [Cl^−^]_i_ in HLF. As shown in figure 3B, 48-hr treatment of HLF with TGF-β resulted in a significant increase in steady state ^36^Cl⁻ accumulation in HLF. Further, T16A_inh_-A01 significantly suppressed both basal and TGF-β-induced steady state ^36^Cl⁻ accumulation in HLF (Fig. 3B). We then used siRNA to knock down ANO1 in HLF. TGF-β increased ^36^Cl⁻ accumulation in cells treated with control siRNA; and, similar to T16Ainh-A01, ANO1 knockdown completely blunted TGF-β-induced accumulation of intracellular ^36^Cl⁻ (Fig. 3C). One difference was that unlike T16A_inh_-A01, siANO1 did not affect intracellular ^36^Cl⁻ levels under the basal conditions (Figs. 3B, 3C, see Discussion). Together, these data suggest that ANO1 activity modulates [Cl^−^]_i_ in HLF, and TGF-β-induced increase in intracellular Cl⁻ (^36^Cl⁻) is fully determined by the increased expression of ANO1.

### Role of ANO1 in TGF-β-induced myofibroblast differentiation

Consistent with our prior work and other publications in the field, treatment of HLF with TGF-β for 48 hours resulted in differentiation to myofibroblasts as evidenced by a profound increase in the expression of the extracellular matrix proteins, collagen-1 (Col1A1) and fibronectin (FN), and dramatic increase in myofibroblast marker, smooth muscle α-actin (SMA) (Fig. 4). These changes paralleled newly found 6-7-fold increases in ANO1 expression (Fig. 4). Unexpectedly, pretreatment of HLF with the ANO1 inhibitor T16A_inh_-A01 blocked the TGF-β-induced increase in ANO1 levels (Fig. 4). More importantly, T16A_inh_-A01 blunted the TGF-β-induced upregulation of α-SMA, Col1A1 and FN without affecting levels of the housekeeping β-actin protein. Attenuation of TGF-β-induced ANO1 expression by T16A_inh_-A01 (Fig. 4) may suggest that in addition to pharmacological inhibition, long-term T16A_inh_-A01 treatment also affects the expression of ANO1 via as yet unknown mechanisms. To corroborate the results with pharmacological blockade of ANO1, we used an siRNA approach. Figure 5 shows efficient ANO1 protein knockdown by ANO1 siRNA and significant decrease in TGF-β-induced expression of α-SMA, Col1A1, and FN very similar to the effect of T16A_inh_-A01.

**Figure 4.**
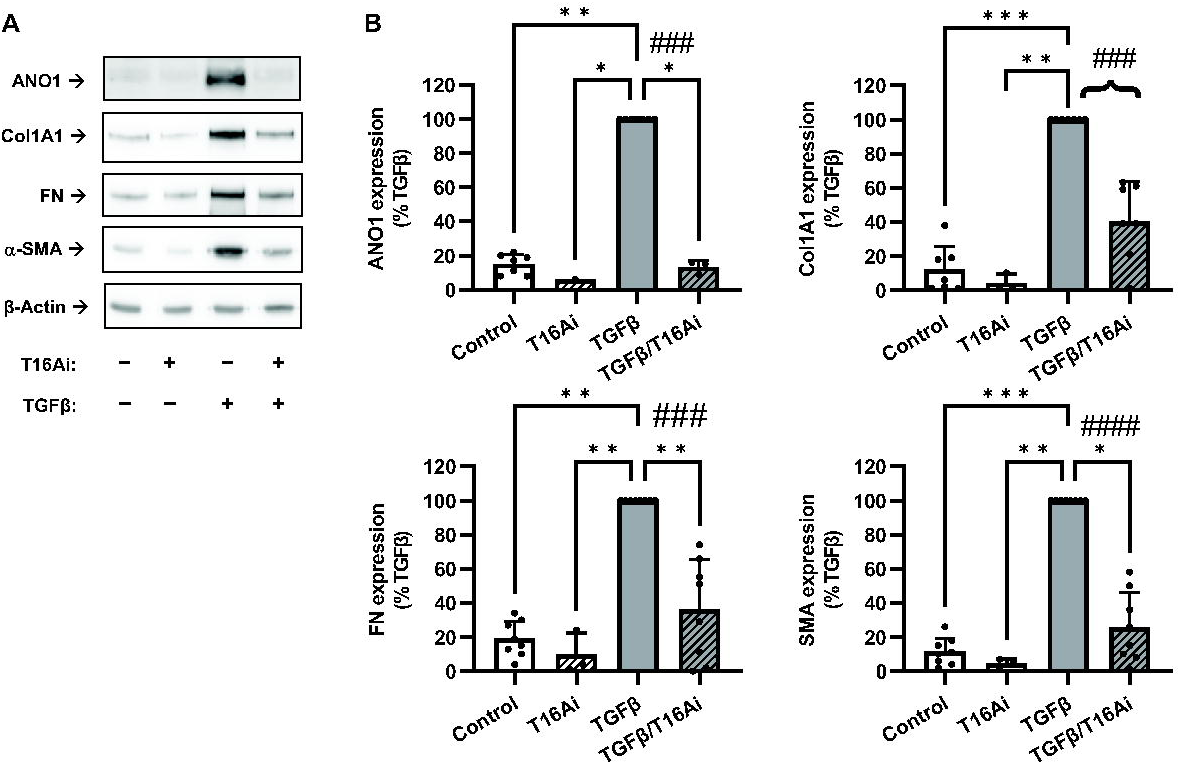
Attenuation of TGF-β-induced myofibroblast differentiation by the ANO1 inhibitor, T16A_inh_-A01. **A,** Representative images of western blot analyses of the effect of the pharmacological inhibitor of ANO1, T16A_inh_-A01 (T16Ai) on myofibroblast differentiation. Serum-starved (48 hr) HLF were pretreated with 30 µM T16Ai or vehicle control for 1hr, followed by treatment with TGF-β (1 ng/ml) for 48 hr. Cell lysates were then probed for immunoreactivity of anactomin-1 (ANO1), collagen 1A1 (Col1A1), fibronectin (FN), and smooth muscle α-actin (SMA). **B,** Quantification of ANO1, Col1A1, FN, and SMA immunoreactivities. Data are mean values ± SD. *p<0.05, **p<0.01, ***p<0.001, vs. TGFβ group, Kruskal-Wallis test with Dunn’s post hoc correction. Brace brackets, ^###^p<0.001; ^####^, p<0.0001, are the results of preplanned comparisons of TGF-β effect with and without ANO1 inhibitor, one-population t-test.

### Control of TGF-β-induced signaling by ANO1 in HLF

We next began exploring the mechanism through which ANO1 controls TGF-β-induced myofibroblast differentiation, starting from established fibrotic pathways. Phosphorylation of Smad2/3 transcription factors by TGF-β receptors is the proximal event in TGF-β signaling. As shown in representative images in Figure 6 (P-Smad2 blots), 30-minute treatment of HLF with TGF-β resulted in phosphorylation of Smad2 without affecting total levels of Smad2. Pretreatment of HLF with the ANO1 inhibitor T16A_inh_-A01 had no effect on TGF-β-induced Smad2 phosphorylation (Fig. 6A), suggesting that the initial signaling event is not affected. We next focused on the downstream, delayed signaling processes that are required for myofibroblast differentiation in response to TGF-β. We and others have previously established the critical role of the RhoA pathway in TGF-β-induced myofibroblast differentiation (19-21). We assessed changes in this pathway by measuring the phosphorylation state of myosin light chain (MLC), which is controlled by RhoA-mediated signaling (37) and represents a widely used indirect assay for RhoA pathway activation (38, 39). Treatment of HLF with TGF-β for 48 hrs increased MLC phosphorylation more than 5-fold, and this effect was significantly inhibited by the pharmacological ANO1 inhibitor, T16A_inh_-A01, whereas the total MLC levels remained unchanged (Fig. 6A, 6B). We also examined the role of ANO1 in the control of AKT phosphorylation in HLF, as has been reported in other cells (40, 41) albeit not in the context of TGF-β. Treatment of HLF with TGF-β for 48 hours resulted in a significant AKT phosphorylation; and this effect was inhibited by T16A_inh_-A01 (Fig. 6A, 6B). To corroborate the data obtained with the pharmacological ANO1 inhibitor, we used the ANO1 knockdown approach. Similarly to T16A_inh_-A01, ANO1 siRNA had no significant effect on Smad2 phosphorylation, but it completely abolished TGF-β-induced phosphorylation of MLC and AKT (Fig. 6C, 6D).

### Control of TGF-β-induced myofibroblast differentiation and signaling by “with-no-lysine (K)” kinase, WNK1

WNK kinases control electrolyte homeostasis and cell volume in renal epithelial cells (42). The WNK1 isoform is ubiquitously expressed, but its function outside of kidney remains poorly understood. It is well established that the kinase activity of WNK1 is activated in response to lowering [Cl^−^]_i_ and conversely inhibited by increased concentrations of intracellular chloride (43). On the other hand, it was recently reported that increased [Cl^-^] promotes a scaffolding (kinase activity-independent) function of WNK1 by bringing together the “mammalian target of rapamycin complex-2” (mTORC2) with its targets, such as Serum/Glucocorticoid Regulated Kinase 1 (SGK1) (44). Therefore, we assessed the role of WNK1 and its kinase activity in TGF-β-induced myofibroblast differentiation and signaling. As shown in Figure 7, siRNA knockdown of WNK1 significantly reduced TGF-β-induced myofibroblast differentiation (Col1A1, FN and SMA expression) as well as RhoA (P-MLC) and AKT (P-AKT) activation. In striking contrast, the pan-WNK kinase inhibitor, WNK463, was largely ineffective at concentrations (10 μM) that are far above its nanomolar WNK1-kinase inhibitory IC50 (45). Together, these data suggest that WNK1 mediates TGF-β-induced myofibroblast differentiation and RhoA and AKT signaling in a kinase-independent manner.

### Control of TGF-β-induced myofibroblast differentiation and signaling by ANO1 and WNK1 in IPF-HLF

We further examined the generality of our findings using lung fibroblasts cultured from IPF lung (IPF-HLF), comparing side-by-side the effects of si*WNK1* and si*ANO1*. As shown in Figure 8, ANO1 was readily detectable under basal conditions in IPF-HLF, and its expression was further induced by TGF-β. Both si*WNK1* and si*ANO1* strongly inhibited myofibroblast differentiation (expression of Col1A1, FN and SMA) as well as two relevant signaling pathways (P-MLC, P-AKT). Interestingly, WNK1 knockdown also attenuated the expression of ANO1 in response to TGF-β (Fig. 8). This suggests a potentially novel feed-forward mechanism in TGF-β signaling through WNK1-dependent upregulation of ANO1.

## DISCUSSION

This study describes a novel and unexpected mechanism of TGF-β-induced myofibroblast differentiation involving the Ca^2+^-activated Cl⁻ channel anoctamin-1 (ANO-1). Our principal findings suggest that ANO1 and the chloride-sensitive signaling protein WNK1 play critical roles in the myofibroblast differentiation process.

We discovered that ANO1 is dramatically upregulated during TGF-β-induced human lung myofibroblast differentiation *in vitro* (Fig. 1). The pathological significance of this discovery is supported by our observation of ANO1 immunoreactivity in human IPF lungs, where it is expressed in the same areas as SMA-positive myofibroblasts (Fig. 2). These findings are novel and have no precedent in the literature. We have previously extensively reviewed the published data on the regulation of ANO1 expression under various conditions and in a variety of cell types (46). ANO1 can be upregulated by several cytokines from the interleukin family (e.g., IL-4, IL-6, IL-13), lysophosphatidic acid, and epidermal growth factor in various cell types. There were several controversial reports on the bidirectional (up or down) regulation of ANO1 expression by serum supplementation or by the G-protein coupled receptor agonist, angiotensin II (reviewed in (46)). In this context, the generality of effect of TGF-β on ANO1 expression we identified is yet to be tested in other cell types. The downstream mechanism(s) of TGF-β-induced ANO1 expression in HLF and other cells also requires further exploration. Nevertheless, a number of important deductions can be made based on the existing literature and findings of the present work (see below).

The most important discovery of this study is that ANO1 expression is a critical, previously unknown step in myofibroblast differentiation. Either pharmacological inhibition or siRNA-mediated knockdown of ANO1 significantly attenuate TGF-β-induced myofibroblast differentiation of HLF (Figs. 4, 5, respectively). This conclusion about ANO1 involvement in myofibroblast transformation is supported by the analysis of expression of three independent myofibroblast markers including the extracellular matrix proteins collagen 1A1 and fibronectin, and the myofibroblast marker smooth muscle α-actin.

Because the most recognized role for ANO1 is in regulation of chloride transport and homeostasis, we first explored the effect of TGF-β on intracellular Cl⁻ levels ([Cl⁻]_i_) and found that this cytokine drives an increase in [Cl⁻]_i_ in HLF, in a manner dependent on ANO1 activity and expression. The ANO1 blocker, T16A_inh_-A01, strongly reduces intracellular Cl⁻ levels under both basal conditions and in the TGF-β-treated HLF (Fig. 3B). ANO1 siRNA prevents the effect TGF-β on the intracellular [Cl^⁻^] but does not modify baseline [Cl⁻] on its own (Fig. 3C). This partial discrepancy in the effects of ANO1 manipulation on baseline [Cl⁻]_i_ may be explained by a) either limited specificity of T16Ainh-A01, or b) by the differential duration of ANO1 inhibition (acute with T16Ainh-A01 vs. chronic with siRNA). T16Ainh-A01 has off-target effects on the ubiquitously expressed LRRC8 volume-regulated anion channels, with the IC_50_ value of ∼5 µM (47). It is also possible that acute inhibition of anion channels (ANO1 and/or LRRC8) can reduce intracellular levels of the negatively charged Cl⁻ via hyperpolarization of HLF. If so, the TGF-β-induced cell transformation would be mediated by changes in membrane potential rather than [Cl⁻]_i_. To address this uncertainty and further test [Cl⁻]_i_ dependency, we examined the long-term effects of DIDS and bumetanide, two inhibitors of electroneutral transporters that reduced [Cl⁻]_i_ (Fig. 3A), on myofibroblast differentiation. As shown in Supplemental Figure S2 (https://doi.org/10.6084/m9.figshare.24514876), bumetanide + DIDS significantly attenuated the effect of TGF-β on FN and SMA expression in HLF. Paradoxically, bumetanide + DIDS increased basal Col1A1 protein expression and did not inhibit, but rather potentiated, the effect of TGF-β on levels of this protein (Supplemental Fig. S2, https://doi.org/10.6084/m9.figshare.24514876). This may point to a non-specific long-term effect of ion transport blockers on HLF cell physiology and deposition of extracellular proteins. However, with the exception of the surprising upregulation of Col1A1 by bumetanide +DIDS, the majority of the experiments employing ANO1 inhibitors (Fig. 4), ANO1 siRNA (Fig. 5), and [Cl⁻]_i_ manipulation (Supplemental Fig. S2, https://doi.org/10.6084/m9.figshare.24514876) are consistent with the idea of [Cl⁻]_i_-dependent regulation of myofibroblast differentiation.

**Figure 5.**
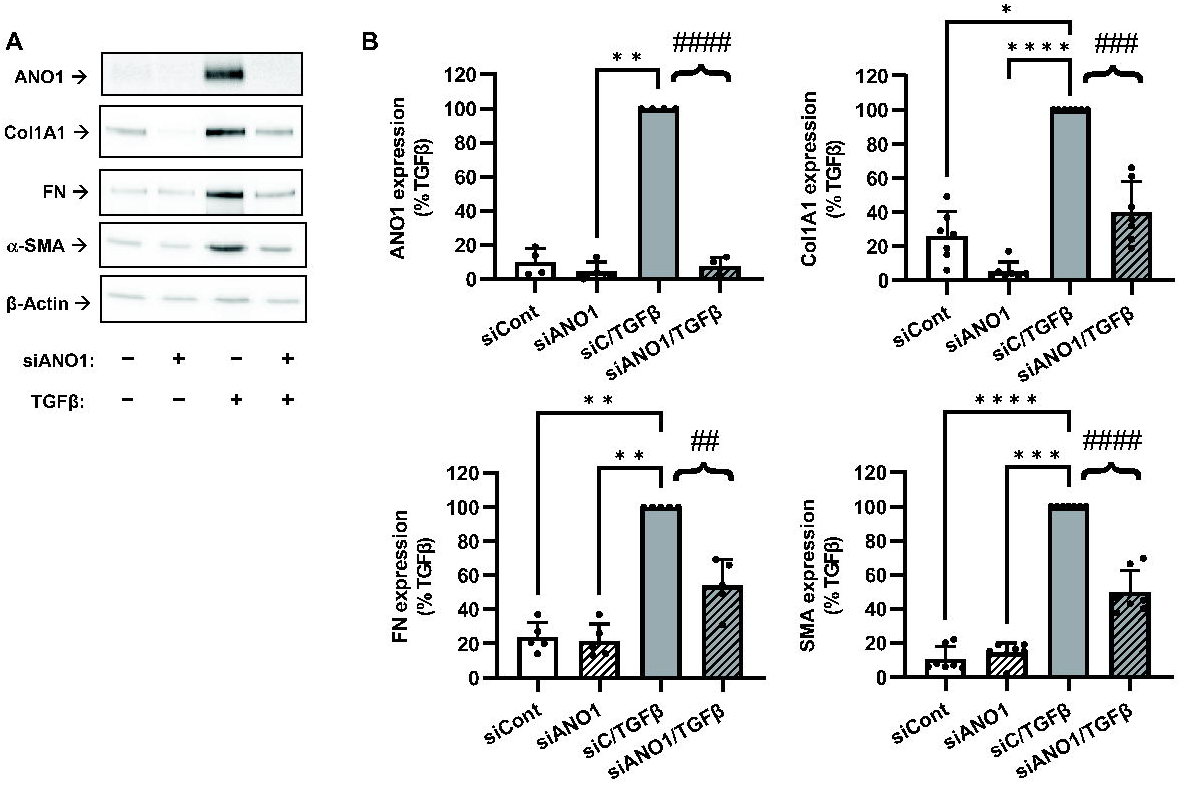
Inhibition of TGF-β-induced myofibroblast differentiation by ANO1 knockdown. **A,** Representative images of western blot analyses of the effect of siRNA targeting ANO1 (siANO1) on myofibroblast differentiation. HLF were transfected with siRNA targeting ANO1 (siANO1) or control RNA (siCont) for 24 hrs, serum starved for 48 hr, and treated with TGF-β or vehicle control for 48 hrs. Cell lysates were then probed for immunoreactivity of anactomin-1 (ANO1), collagen 1A1 (Col1A1), fibronectin (FN), and smooth muscle α-actin (SMA). **B,** Quantification of ANO1, Col1A1, FN, and SMA immunoreactivity. Data are mean values ±SD. *p<0.05, **p<0.01, ***p<0.001, vs. TGFβ group, Kruskal-Wallis test with Dunn’s post hoc correction. Brace brackets, ^##^p<0.01 ^###^p<0.001; ^####^, p<0.0001, are the results of preplanned comparisons of TGF-β effect in cells treated with either control siRNA (siC) or siANO1, one-population t-test.

The effect of TGF-β on ANO1 expression needs to be considered in the context of known mechanisms driving TGF-β-dependent changes in gene expression. (i) ANO1 can be a direct target of canonical SMAD signaling, and the ANO1 promoter may have previously unrecognized SMAD-binding elements. (ii) ANO1 levels may be indirectly modulated by Smad-dependent gene transcription through regulation of downstream signaling pathways. For example, ANO1 expression is induced by IL-6 (through STAT3 transcription factor) (48) while IL-6 expression is induced by TGF-β also by recruiting the STAT3 pathway (49, 50); (iii) ANO1 transcription may be under control of myocardin/serum response factor (SRF) as seen in vascular smooth muscle cells (51). Given our previous findings that TGF-β promotes the SMAD-dependent activation of the RhoA/SRF pathway in HLF (20, 21), the TGF-β/SRF-dependent transcription of ANO1 is also plausible.

Findings presented in Fig. 6 indicate that inhibition of ANO1 does not affect the immediate TGF-β signaling involving Smad2 phosphorylation, but does block downstream signaling events, including phosphorylation of MLC and AKT (Fig. 6). The net phosphorylation of MLC (controlled by RhoA-dependent inactivation of MLC phosphatase (37)) has been commonly used for an indirect assessment of RhoA activation (38, 39). Our previous work established a critical role of the RhoA/SRF pathway in TGF-β-induced myofibroblast differentiation (19-21). The present study found complete blockade of MLC phosphorylation by pharmacological inhibition of ANO1 or ANO1 siRNA knockdown (Fig. 6). Based on these observations, we argue that TGF-β-dependent activation of the RhoA/SRF pathway could be one of the mechanisms by which ANO1 controls myofibroblast differentiation. On the other hand, ANO1 transcription has been reported to be driven by SRF (51), potentially suggesting the reciprocal regulation and a positive feed-forward loop mechanism involving both SRF and ANO1. Besides the RhoA/SRF axis, we also found that inhibition and siRNA knockdown of ANO1 both reduce TGF-β-dependent phosphorylation of AKT (Fig. 6). A role of ANO1 in regulation of AKT phosphorylation has been reported in other cells (40, 41), but not in the context of TGF-β stimulation. AKT plays an important role in TGF-β-induced myofibroblast differentiation (24) and may additionally contribute to the pro-fibrotic effects of ANO1.

**Figure 6.**
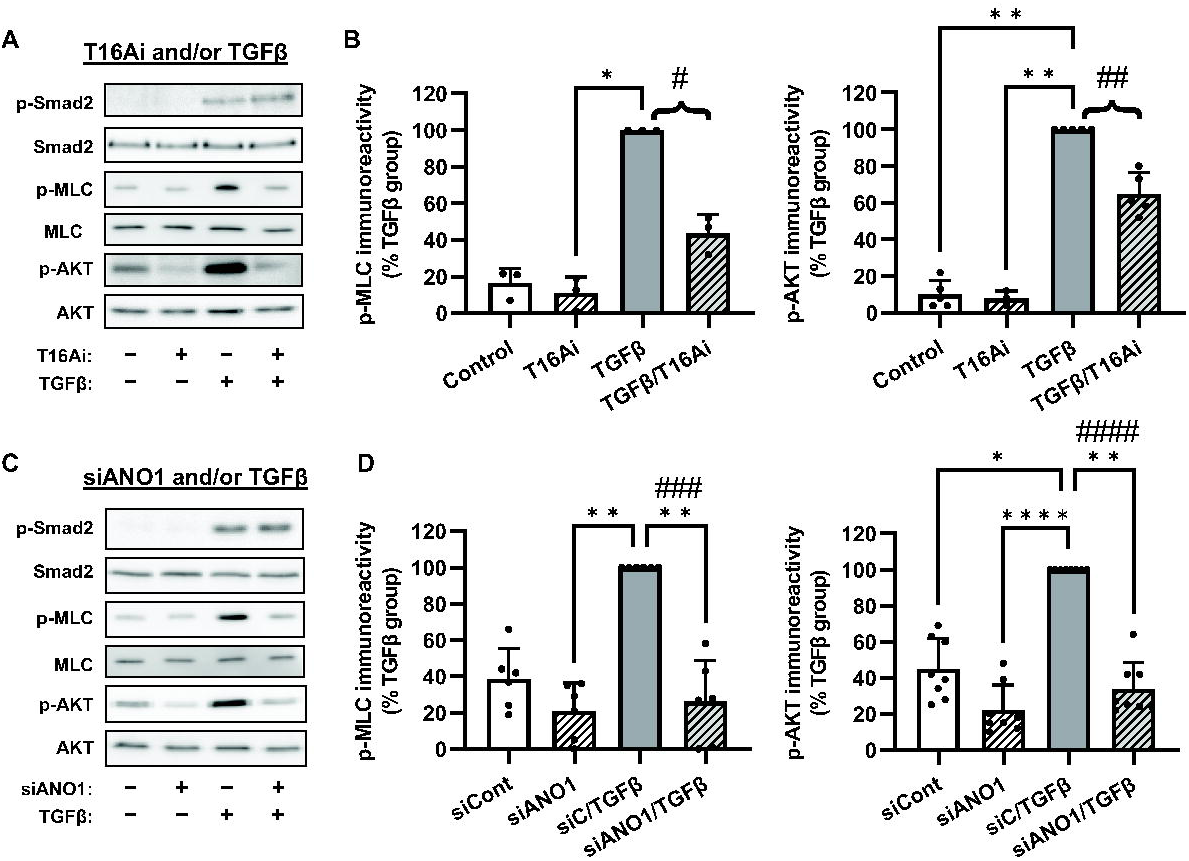
ANO1 contributes to TGF-β-induced RhoA and AKT signaling without affecting Smad2 phosphorylation. **A,** Representative images of western blot analyses of the pharmacological effect of the ANO1 inhibitor, T16Ainh-A01 (T16Ai), on intracellular signaling pathways in HLF. Serum-starved (48 hr) cells were pretreated for 1 hr with T16Ai, followed by treatment with TGF-β (1 ng/ml) for either 30 min (p-Smad2, Smad2), or 48 hr (p-MLC, MLC, p-AKT, AKT). **B,** Quantification of p-MLC and p-AKT immunoreactivities in T16Ai-treated cells. Data represent the mean immunoreactivity values ±SD normalized to TGF-β group within the same experiment. *p<0.05, **p<0.01, vs. TGF-β group, Kruskal-Wallis test with Dunn’s post hoc correction. Brace brackets, #p<0.05; ##, p<0.01, are the results of preplanned comparisons of the TGFβ effect in cells treated with or without T16Ai, one population t-test. **C,** Representative images of western blot analyses of the effects of the siRNA targeting ANO1 (siANO1) on intracellular signaling pathways in HLF. HLF cells were transfected with the siRNA targeting ANO1 (siANO1) or control RNA (siCont) for 24 hrs, serum starved for 48 hrs, and treated with TGFβ (1 ng/ml) for either 30 min (p-Smad2, Smad2), or for 48 hr (p-MLC, MLC, p-AKT, AKT). **D,** Quantification of p-MLC and p-AKT immunoreactivities in siCont and siANO1-treated cells. Data represent mean immunoreactivity values ±SD normalized to β-actin loading control and TGF-β group within the same experiment. * *p<0.05, **p<0.01, ***p<0.001, ****p<0.0001, vs. TGF-β group, Kruskal-Wallis test with Dunn’s post hoc correction. Brace brackets, ###p<0.001; ####, p<0.0001, are the results of preplanned comparisons of the TGF-β effect in cells treated with control siRNA (siC) or siANO1, one population t-test.

What is a common denominator among intracellular Cl⁻ levels, TGF-β signaling, and ANO1 expression? Our study uncovers a novel, Cl⁻-dependent, mechanism for TGF-β-induced myofibroblast differentiation and RhoA and AKT signaling that involves the chloride-sensing protein kinase WNK1. All the effects of TGF-β we observed on myofibroblast differentiation and signaling were inhibited by WNK1 knockdown (Fig. 7). Yet, TGF-β signaling and induction of myofibroblast markers were largely insensitive to the actions of the pan-WNK kinase inhibitor, WNK463 (Fig. 7). These new findings are consistent with a recently reported kinase-independent role of WNK1 in activation of mTORC2 signaling under conditions of high intracellular [Cl^-^] (44). According to that study, binding of Cl⁻ ions to the autoinhibitory domain of WNK1 blocks its kinase activity but promotes WNK1 association with the mTOR complex 2, followed by activation of downstream signaling molecules (44). It is established that mTORC2 promotes AKT autophosphorylation at serine-473 (52), the phosphorylation site we assessed in this study (Figs. 6-8). mTORC2 was also shown to be required for RhoA activity in rat uterine leiomyoma cells and smooth muscle-like lymphangioleiomyomatosis (LAM) cells (53), in mesenchymal stem cells (54), and in neutrophils (55).

**Figure 7.**
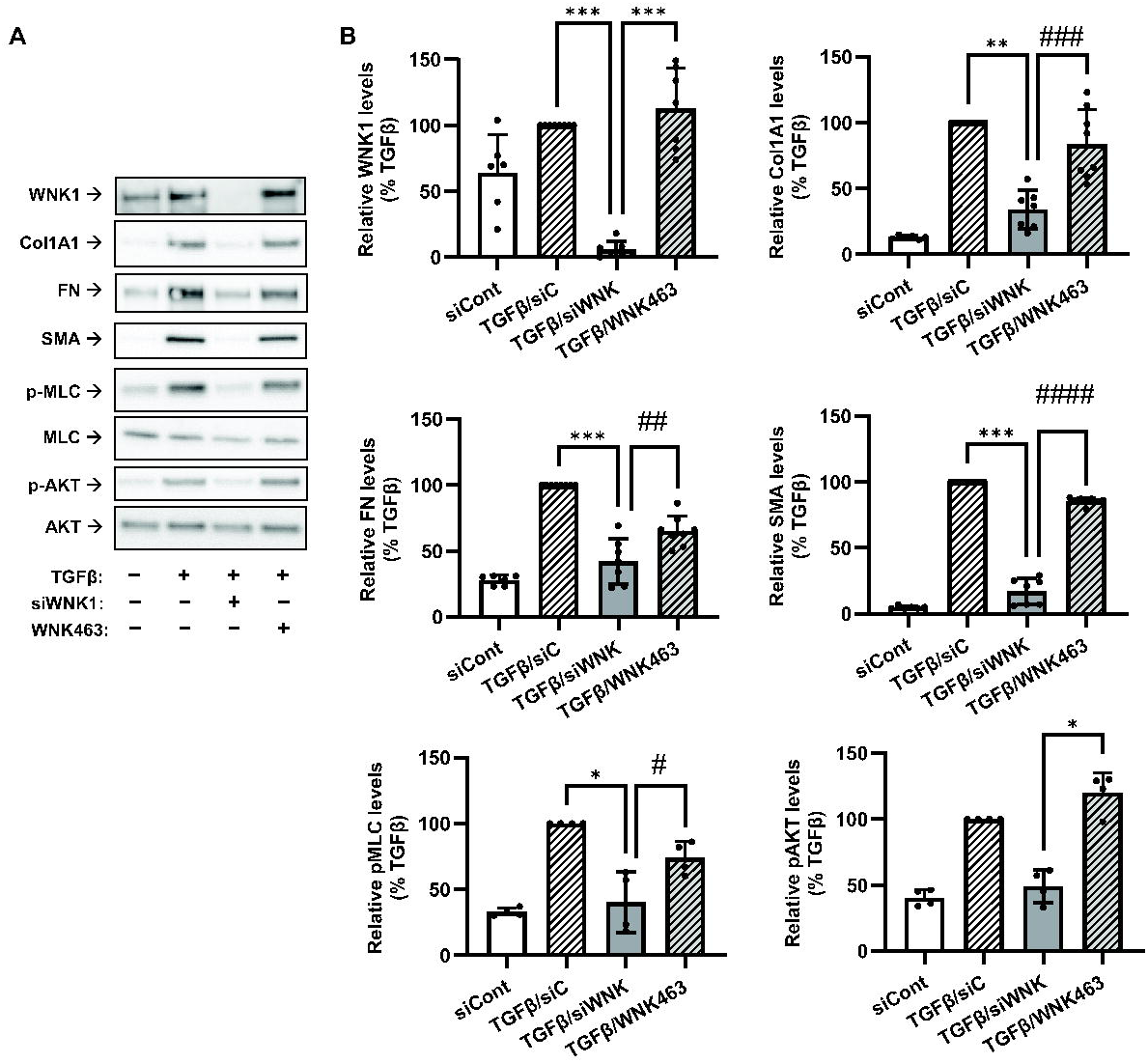
WNK1 mediates TGFβ-induced myofibroblast differentiation in a kinase-independent manner. **A,** Representative images of western blot analyses of the effects of siRNA targeting WNK1 (siWNK1) and the WNK kinase inhibitor WNK463 on myofibroblast markers and intracellular signaling pathways in HLF. HLF were transfected with siRNA targeting ANO1 (siANO1) or control RNA (siCont) for 24 hrs, serum starved for 48 hrs, pretreated with or without WNK1 inhibitor WNK463 (10 µM) for 1 hr, followed by treatment with TGF-β (1 ng/ml) or vehicle control for 48 hrs. Cell lysates were then probed for the immunoreactivity of WNK1, collagen 1A1 (Col1A1), fibronectin (FN), smooth muscle α-actin (SMA), p-MLC, MLC, p-AKT, and AKT. **B,** Quantification of the immunoreactivity of WNK, Col1A1, FN, SMA, p-MLC, and p-AKT. Data are mean IR values ±SD normalized to β-actin loading control and TGFβ group within the same experiment. *p<0.05, **p<0.01, ***p<0.001, vs. TGF-β group, Kruskal-Wallis test with Dunn’s post hoc correction. Brace brackets, #p<0.05; ##, p<0.01, are the results of preplanned comparisons of the TGFβ effect in cells treated with siWNK1 or the WNK1 inhibitor WNK463, t-test.

**Figure 8.**
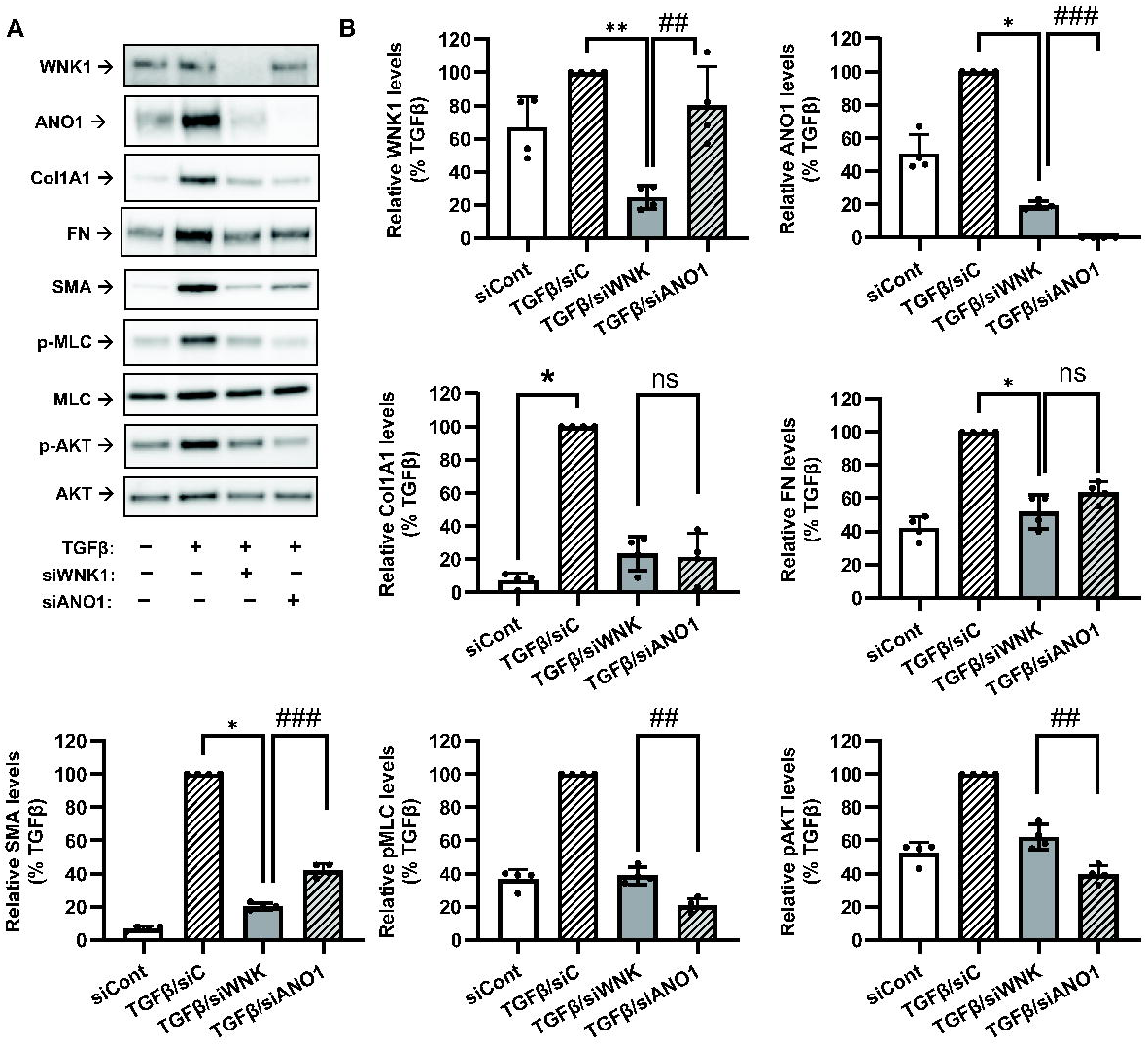
Relative contributions of WNK1 and ANO1 to TGFβ-induced myofibroblast differentiation and signaling in human lung fibroblasts isolated from the IPF lungs (IPF-HLF). **A,** Representative images of western blot analyses of the effects of siRNA targeting WNK1 (siWNK1) or ANO1 (siANO1) on myofibroblast differentiation and intracellular signaling in IPF-HLF. Cells were transfected with siRNA targeting WNK1 (siWNK1), ANO1 (siANO1) or control RNA (siCont) for 24 hrs, serum starved for 48 hrs, followed by treatment with TGF-β (1 ng/ml) or vehicle control for 48 hrs. Cell lysates were then probed for the immunoreactivity of WNK1, collagen 1A1 (Col1A1), fibronectin (FN), smooth muscle α-actin (SMA), p-MLC, MLC, p-AKT, and AKT. **B,** Quantification of the immunoreactivity of WNK, Col1A1, FN, SMA, p-MLC, and p-AKT. Data are mean values ±SD normalized to TGF-β group within the same experiment. *p<0.05, **p<0.01, vs. TGF-β group, Kruskal-Wallis test with Dunn’s post hoc correction. Brace brackets, #p<0.05; ##, p<0.01, are the results of preplanned comparisons of the TGF-β effect in cells treated with or without T16Ai, one population t-test.

In a separate noteworthy series of experiments, we explored the relative contributions of ANO1 and WNK1 to regulation of myofibroblast markers in HLF that were isolated from IPF lungs (IPF-HLF). In IPF fibroblasts, ANO1 expression is readily detectable even in the absence of TGF-β (Fig. 8). Treatment with TGF-β causes additional increases in ANO1 levels, upregulates expression levels of the myofibroblast markers Col1A1, FN, and SMA, and stimulates phosphorylation of MLC and AKT (Fig. 8). Consistent with findings in non-IPF fibroblasts (Figs. 4-7), the effects of TGF-β are strongly reduced or completely inhibited by siRNA targeting either ANO1 or WNK1 (Fig. 8). However, we also found that downregulation of WNK1 reduces the levels of the “upstream” ANO1 (Fig. 8). This latter discovery points to the bi-directional positive interaction between ANO1 and WNK1, which may constitute a feed-forward mechanism in myofibroblast differentiation.

In conclusion, we propose a **novel model for myofibroblast differentiation**, which is summarized in Fig. 9. Our study uncovered an entirely unexpected role of intracellular Cl⁻ in pro-fibrotic TGF-β signaling. The new hypothetical mechanism incorporates the following signaling steps. (i) TGF-β stimulates expression of ANO1, resulting in an increase in intracellular [Cl⁻] in HLF. (ii) High intracellular chloride promotes a scaffolding, kinase-independent function of WNK1 and drives its association with mTORC2 and activation of downstream signaling pathways (44). (iii) mTORC2 activates AKT and RhoA, all of which drive myofibroblast differentiation (19-21, 24, 56, 57). P-MLC, activated by RhoA, also promotes myofibroblast differentiation (58, 59) and fibronectin assembly (60) through myosin-dependent cell contraction. (iv) Interestingly, ANO1 expression is both a prerequisite for the TGF-β WNK1/mTORC2 signaling and is also dependent on WNK1 expression. This suggests a bidirectional feed-forward mechanism contributing to myofibroblast differentiation. The validity of this model and its relevance to diverse aspects of myofibroblast pathophysiology deserves further exploration, particularly in *in vivo* models of pulmonary fibrosis.

**Fig. 9.**
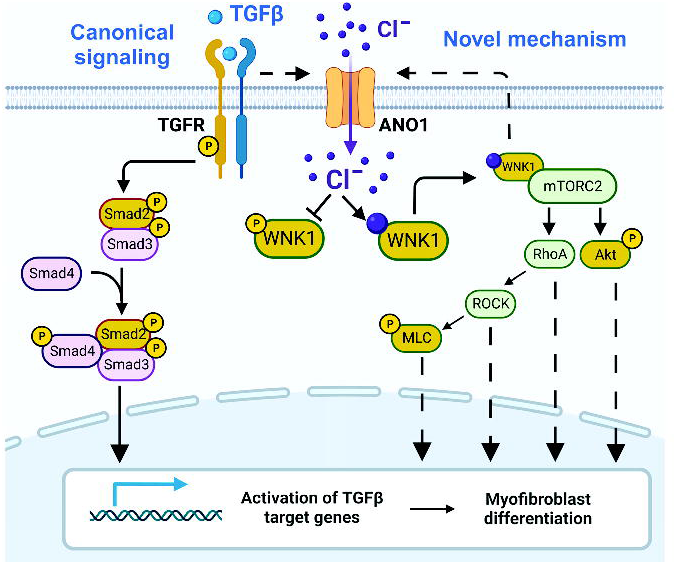
Novel model for TGF-β-induced myofibroblast differentiation mediated by ANO1 and WNK1. The left part of the diagram illustrates the canonical model of the TGF-β-induced myofibroblast differentiation involving Smad signaling. The right part of the diagram depicts novel mechanism which this study suggests. (i) TGF-β upregulates the expression of ANO1 and elevates intracellular [Cl□]. (ii) Increased [Cl□]i stabilizes autoinhibitory conformation of WNK1 and prompts its association with and activation of mTORC2. (iii) mTORC2 activates AKT and RhoA pathways. (iv) AKT and RhoA promote myofibroblast differentiation via downstream signaling mechanisms, also including P-MLC. (v) WNK1 signaling forms a feedforward mechanism for ANO expression. Yellow color represents signaling molecules whose contribution was tested in this study. Prepared with BioRender.

## ACKNOWLEDGEMENTS

The Authors thank J.W. Nalwalk for critical reading and helpful suggestions on the manuscript. Graphical summary in Fig. 9 was created with BioRender (BioRender.com).

## AUTHOR CONTRIBUTIONS

Eleanor B. Reed: conceived and designed research; performed experiments, interpreted results of experiments, analyzed data, approved final version of manuscript.

Shaina Orbeta: performed experiments, analyzed data, interpreted results of experiments, approved final version of manuscript.

Bernadette A. Miao: conceived and designed research, performed experiments, analyzed data, interpreted results of experiments, prepared figures, edited and revised manuscript, approved final version of manuscript.

Albert Sitikov: performed experiments, analyzed data, interpreted results of experiments, prepared figures, approved final version of manuscript.

Bohao Chen: performed experiments, analyzed data, interpreted results of experiments, approved final version of manuscript.

Irena Levitan: conceived and designed research, performed experiments, analyzed data, interpreted results of experiments, approved final version of manuscript.

Julian Solway: interpreted results of experiments, edited and revised manuscript, approved final version of manuscript.

Gökhan M. Mutlu: interpreted results of experiments, edited and revised manuscript, approved final version of manuscript.

Yun Fang: interpreted results of experiments, edited and revised manuscript, approved final version of manuscript.

Alexander A. Mongin: conceived and designed research, performed experiments, analyzed data, interpreted results of experiments, prepared figures, drafted manuscript, edited and revised manuscript, approved final version of manuscript.

Nickolai O. Dulin: conceived and designed research, performed experiments, analyzed data, interpreted results of experiments, prepared figures, drafted manuscript, edited and revised manuscript, approved final version of manuscript.

