## Supplemental Figure S1 for "Anoctamin-1 is induced by TGF-beta and contributes to lung myofibroblast differentiation"

Supplemental Figure 1 • E.B. Reed *et al.*

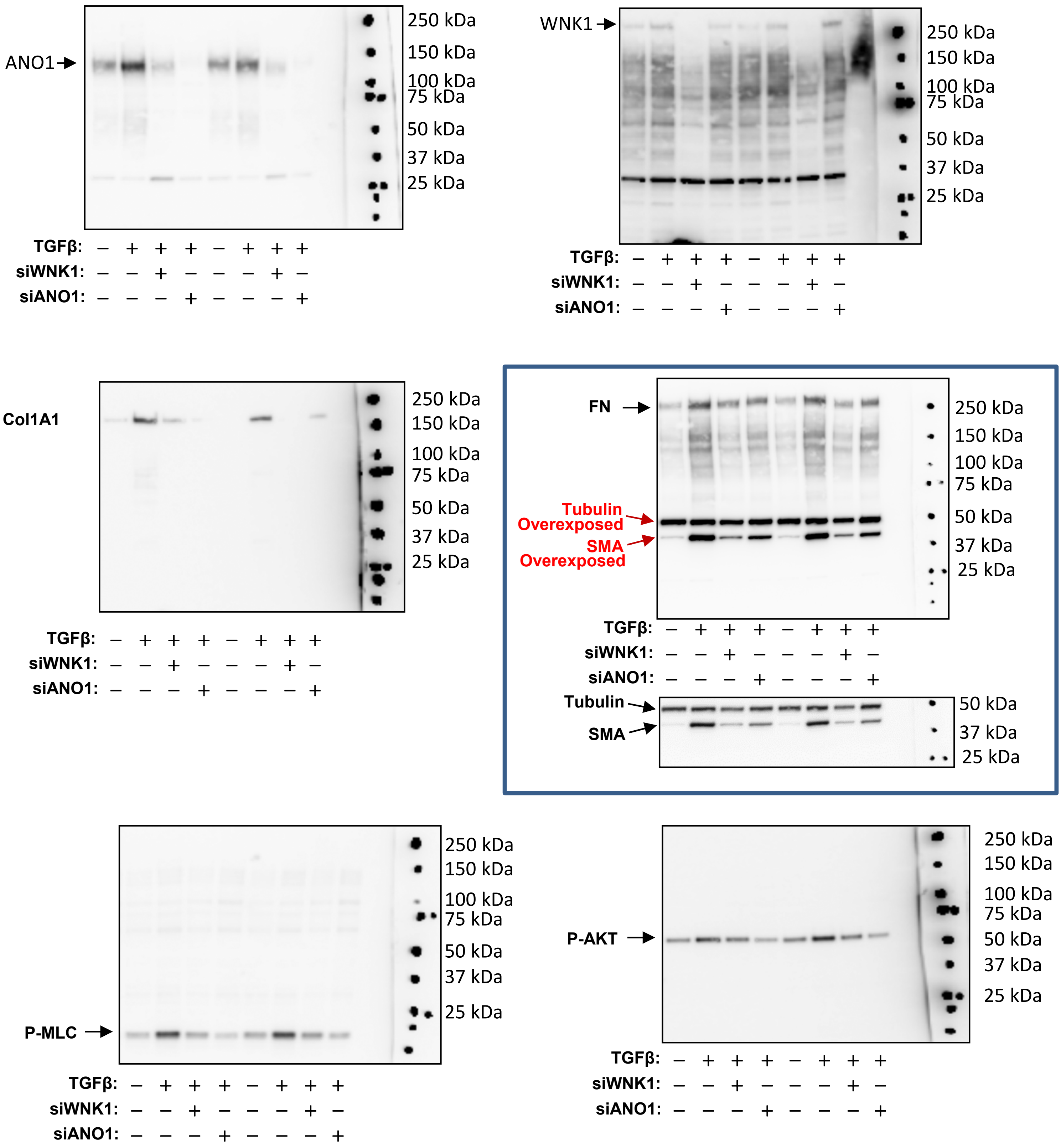


**Representative full-length blots for ANO1, WNK1, Col1A1, FN, SMA, Tubulin, P-MLC and P-AKT**

Because molecular weights of FN, SMA and Tubulin do not overlap, the mixture of corresponding primary antibodies was used and ECL images were obtained under different exposures below the saturation levels for each proteins. The separate experiments confirmed that SMA or Tubulin antibodies recognize single corresponding bands, and there is no detectable FN smear at or below 50kDa (data not shown).
