## Supplemental Figure S2 for "Anoctamin-1 is induced by TGF-beta and contributes to lung myofibroblast differentiation"

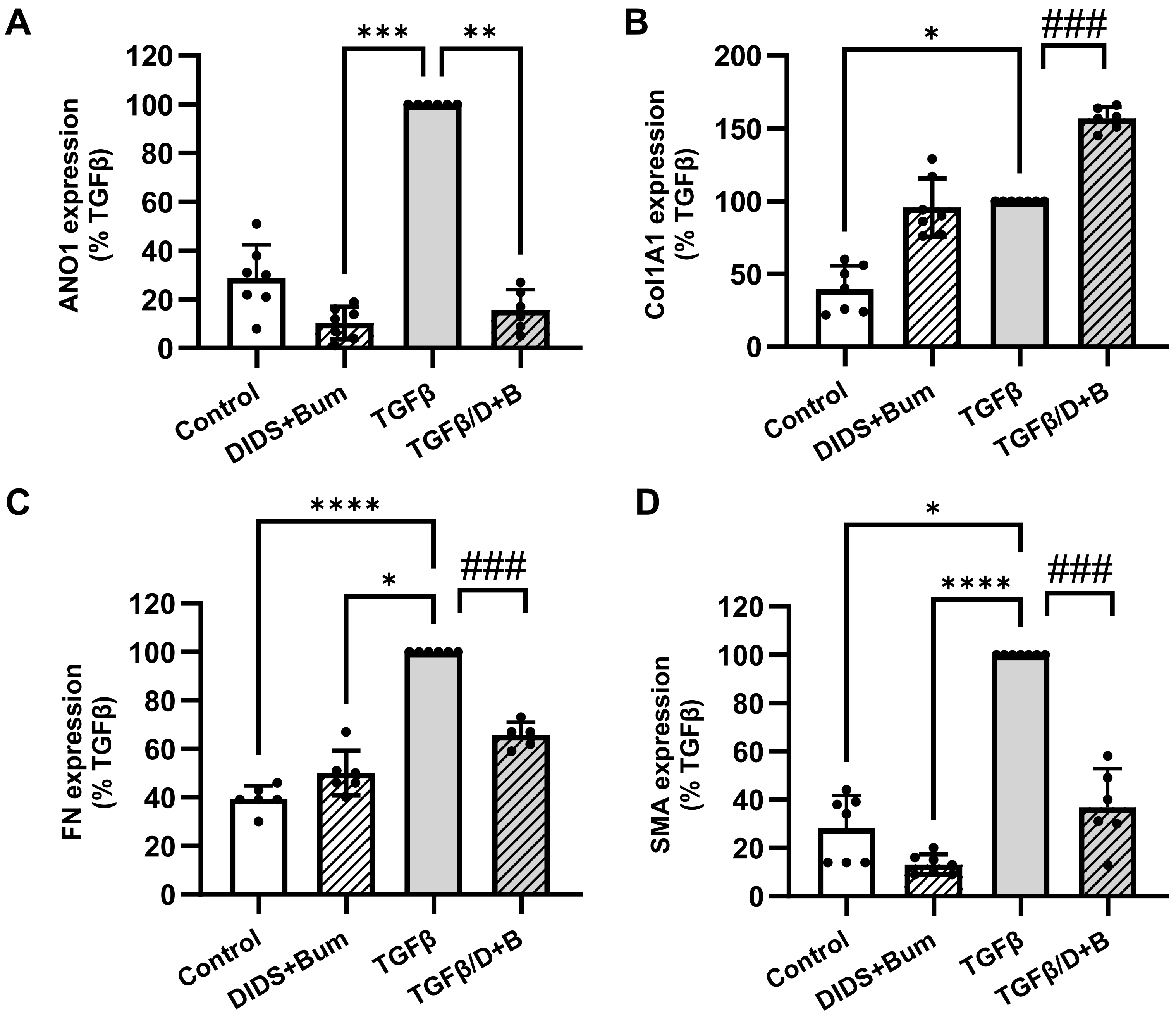


**The effect of long-term treatment with DIDS and bumetanide on ANO1 expression and the TGF-β-induced myofibroblast differentiation.**

Serum-starved (48 hr) HLF were treated with 1 mM DIDS, 20 μM bumetanide and 1 ng / ml TGF-β for 48 hr. Quantification of ANO1 (A), Col1A1 (B), FN (C), and SMA (D) immunoreactivities. Data are mean values ± SD. *p<0.05, **p<0.01, ***p<0.001, vs. TGFβ group, Kruskal-Wallis test with Dunn's post hoc correction. Brace brackets, ###p<0.001, are the results of preplanned comparisons of TGF-β effect with and without DIDS+bumetanide, one-population t-test.
